# Motif signatures of transcribed enhancers

**DOI:** 10.1101/188557

**Authors:** Dimitrios Kleftogiannis, Haitham Ashoor, Nikolaos Zarokanellos, Vladimir B. Bajic

**Affiliations:** Centre for Evolution and Cancer, Division of Molecular Pathology, The Institute of Cancer Research (ICR), London, SW7 3R, United Kingdom; Computational Bioscience Research Center (CBRC), Computer, Electrical and Mathematical Sciences and Engineering Division (CEMSE), King Abdullah University of Science and Technology (KAUST), Thuwal, 23955-6900, Saudi Arabia; Red Sea Research Center (RSRC), Biological and Environmental Sciences & Engineering Division (BESE), King Abdullah University of Science and Technology (KAUST), Thuwal, 23955- 6900, Saudi Arabia

## Abstract

In mammalian cells, transcribed enhancers (TrEn) play important roles in the initiation of gene expression and maintenance of gene expression levels in spatiotemporal manner. One of the most challenging questions in biology today is how the genomic characteristics of enhancers relate to enhancer activities. This is particularly critical, as several recent studies have linked enhancer sequence motifs to specific functional roles. To date, only a limited number of enhancer sequence characteristics have been investigated, leaving space for exploring the enhancers genomic code in a more systematic way. To address this problem, we developed a novel computational method, TELS, aimed at identifying predictive cell type/tissue specific motif signatures. We used TELS to compile a comprehensive catalog of motif signatures for all known TrEn identified by the FANTOM5 consortium across 112 human primary cells and tissues. Our results confirm that distinct cell type/tissue specific motif signatures characterize TrEn. These signatures allow discriminating successfully a) TrEn from random controls, proxy of non-enhancer activity, and b) cell type/tissue specific TrEn from enhancers expressed and transcribed in different cell types/tissues. TELS codes and datasets are publicly available at http://www.cbrc.kaust.edu.sa/TELS.

## INTRODUCTION

In mammalian cells, spatial and temporal activation of gene transcription and maintenance of expression levels is coordinated (mainly) by interactions between DNA regulatory elements, the most prominent being promoters and enhancers^1^. Promoters surround the Transcription Start Site (TSS) of genes and represent the class of proximal regulatory elements. Specific regions in promoters are used as binding sites responsible for recruiting and anchoring the transcriptional machinery^2^. On the other hand, enhancers, frequently called distal regulatory elements, are positioned few or many thousand base pairs (bp) downstream or upstream from the TSSs of genes. Typically enhancers activate their target genes via physical interactions with Transcription Factors (TFs), co-activators and/or via chromatin remodeling processes^3,4^. Results obtained from Cap Analysis of Gene Expression (CAGE) show that transcription in enhancers mediated by Polymerase II (POL2) occurs on a genome-wide scale^5^. Enhancers’ transcription produces eRNAs, a class of non-coding RNAs whose functions are unclear^6,7^. It is interesting to note that it may be difficult to clearly separate, based on transcriptional activation similarity, enhancers and promoters since both act as promoters but generate different classes of transcripts^8-10^.

Several enhancer identification methods, covering both experimental and computational approaches, has been subject of reviews^11,12^. Using the available enhancer-related information^13,14^, a number of studies linked variations in enhancer sequences to disease phenotypes, and development/progression of cancer^15-19^. Thus, deciphering the genomic characteristics of enhancers may help better understanding enhancers’ functional roles.

Up to now, there are several approaches to analyse enhancers’ DNA characteristics and associate sequence properties to enhancers’ activities^20,21^. However, only a limited number of cases in terms of studied enhancer sequences, sequence motifs (e.g., kmers of length 6-8 bp), organisms (e.g., mouse or Drosophila), and tissues, have previously been examined or validated experimentally^22^ (i.e., by Massive Parallel Reporter Assays – MPRAs), leaving space for further investigations.

With all of the above issues in mind, we introduce TELS (Transcribed Enhancer Landscape Search) a novel bioinformatics algorithm that applies logistic regression (LR) coupled with dimensionality reduction techniques to identify systematically the most informative combinations of short sequence motifs of TrEn in the human genome. Our primary aim is to identify combinations of short sequence motifs (equally denoted as DNA signatures or motif signatures) in TrEn sequences that operate in a cell type/tissue specific manner and to investigate the degree to which the identified motifs lead to accurate discrimination of TrEn. As importantly, TELS addresses the problem of discriminating TrEn from different negative controls using exclusively DNA sequence characteristics and without prior knowledge of Transcription Factor Binding sites (TFBSs) from ChIP-seq or Position Weight Matrices (PWMs).

As a case study, we applied TELS to the Atlas of CAGE-defined TrEn from ref. (5) covering 112 human primary cells and tissues. Our analysis identified DNA signatures that are predictive of cell type/tissue specific TrEn. The identified combinations of motifs discriminate successfully TrEn from random controls, proxy of non-enhancer activity, as well as enhancers transcribed exclusively in cell type/tissue specific manner from enhancers transcribed exclusively in different primary cells or tissues. Our results demonstrate that the proposed method leads to the discovery of informative motif signatures, opening possibilities for analysing systematically the genomic landscape of human TrEn.

## RESULTS

### Identification of DNA signatures of TrEn in all FANTOM5 facets

We used TELS to compile a comprehensive catalog of motif signatures that discriminate effectively TrEn in 112 cell types/tissues included in the ‘all facets’ dataset from ‘all facets random controls’ (see Materials and Methods for data description). The average classification performance across 112 cell types/tissues in terms of PPV and GM is 85.94% and 86.06%, respectively (Supplementary Figure 1). Considering a performance threshold of 80% PPV, we observe that the identified motif signatures classify TrEn with high accuracy in 95% of cases (107 out of 112 cell types/tissues). This suggests that the identified combinations of sequence motifs capture a great portion of the sequence specificities required in TrEn.

Figure 1 shows that the performance maximization across different cell types/tissues is achieved using different combinations of motifs. The number of selected motifs ranges from 204 to four, which correspond to the maximum and minimum numbers of motifs that discriminate efficiently TrEn from random controls, a proxy of non-enhancer activity. It is apparent that the model complexity, in terms of number of predictor variables, does not affect the classification efficiency, which appears almost always quite high.

**Figure 1:**
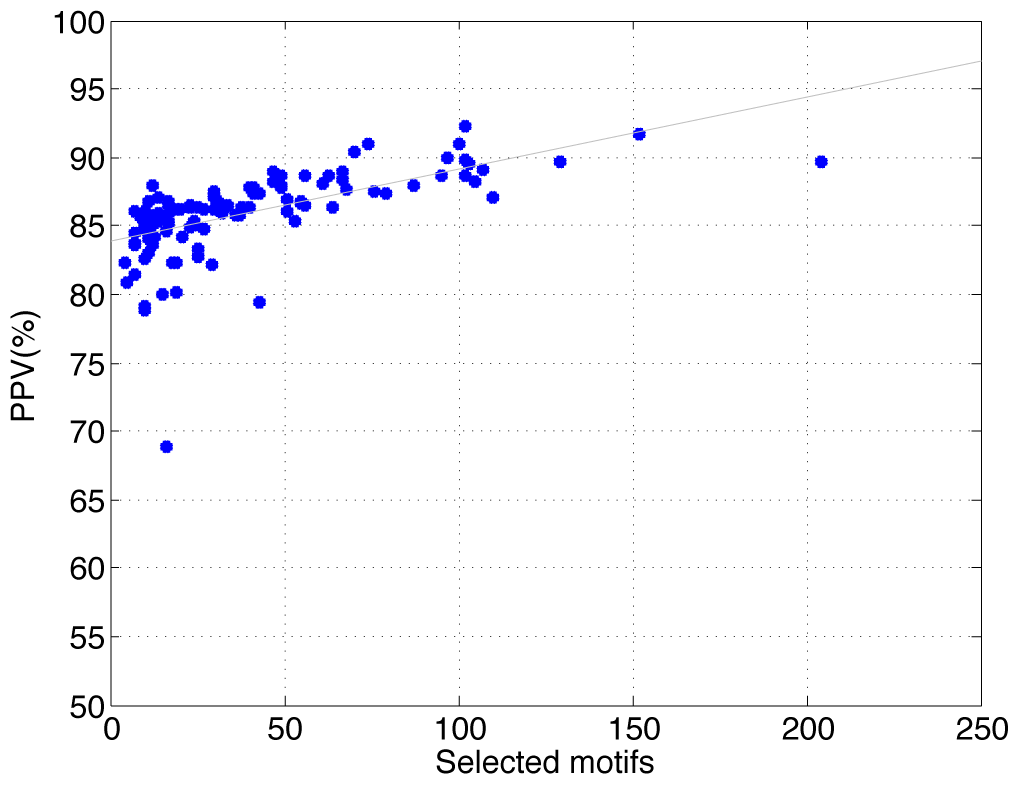
Number of selected motifs across 112 cell types/tissues versus the corresponding PPV (%) for identifying TrEn (‘all-facets’ dataset) from random controls.

To investigate further the identified sets of motif signatures, we select nine tissues that belong to three different developmental stages namely ectoderm (brain, spinal-cord, and eye), mesoderm (kidney, heart and spleen) and endoderm (lung, liver and pancreas) according to the Embryonic Development and stem cell compendium (https://discovery.lifemapsc.com/in-vivodevelopment). For the selected tissues we generate the pairwise similarity matrix using Jaccard index (i.e., considering exact matches) for the sets of motif signatures (Figure 2a). We also generate the pairwise similarity matrix using Jaccard index (i.e., considering exact matches) for the corresponding sets of input TrEn sequences (Figure 2b). Another graphical representation using dendrograms is available in Supplementary Figure 7. Our results show that different sets of TrEn are ‘weakly’ similar to each other based on their input enhancer sequences (Figure 2b). Tissues that belong to ectoderm, mesoderm and endoderm have average pairwise input similarity of 0.129, 0.056 and 0.207, respectively, whereas pancreas and liver appear the most similar tissues based on their input sequences. As expected, tissues that are more similar in terms of input sequences have somewhat more similar sets of motif signatures. However, the level of similarity across the identified sets of informative motifs is quite low (Figure 2a). The average pairwise Jaccard similarity across all nine sets of motif signatures is 0.022. Mesoderm tissues have completely disjoint sets of motifs (i.e., for kidney, heart and spleen the number of selected motifs is only 8, 8 and 4, respectively). Apparently, heart tissue appears similar to lung and pancreas that belong to different developmental stage, both in terms of input sequences and selected motifs.

**Figure 2:**
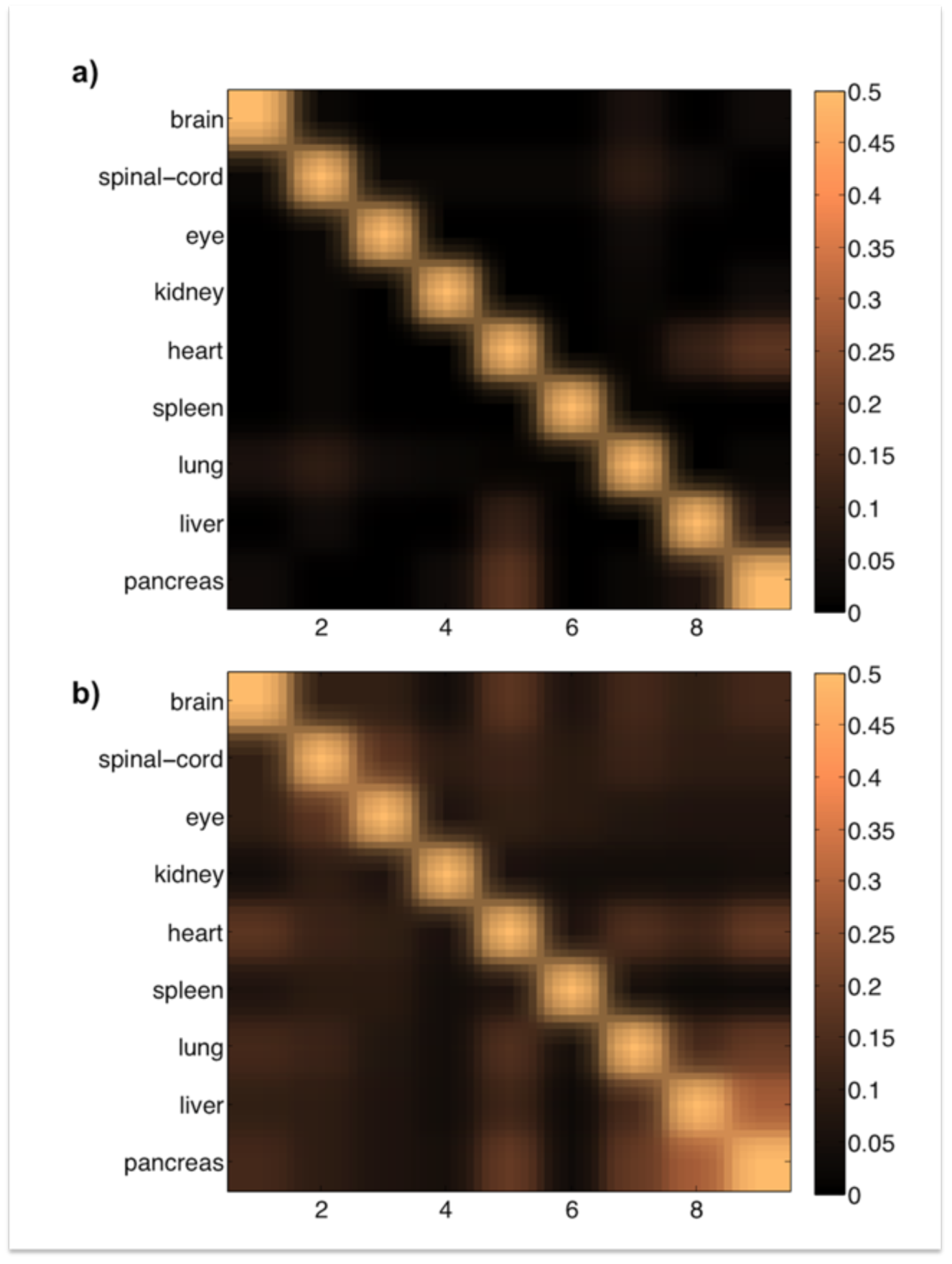
Discriminative identification of TrEn (‘all-facets’ dataset) from random controls using the identified tissue-specific motif signatures: (a) Similarity matrix based on Jaccard index constructed from the selected sets of motif signatures; (b) Similarity matrix based on Jaccard index constructed from the actual input enhancer sequences.

These observations suggest that TrEn display cell type/tissue specific motif signatures that are identified by TELS. Overall, the results presented in this subsection support the hypothesis that specific genomic characteristics enable TrEn to operate in a highly cell type/tissue specific manner and for this reason the identified motifs vary across different cell types/tissues.

### Identification of DNA signatures of TrEn expressed in at least one cell type/tissue from FANTOM5

In this subsection we identify motif signatures that allow us to discriminate the ‘robust set’ of TrEn from the ‘robust set random control’ with maximized classification performance (see Materials and Methods for data description). Our results show that TELS achieves this goal with an average PPV and GM of 79.70% and 80.47%, respectively. Figure 3 shows the characteristic ROC curve, using the most informative combination of 31 motifs. Overall, using this motif signature we report an average Area Under Curve (AUC) of 0.854 with standard deviation of 0.009.

**Figure 3:**
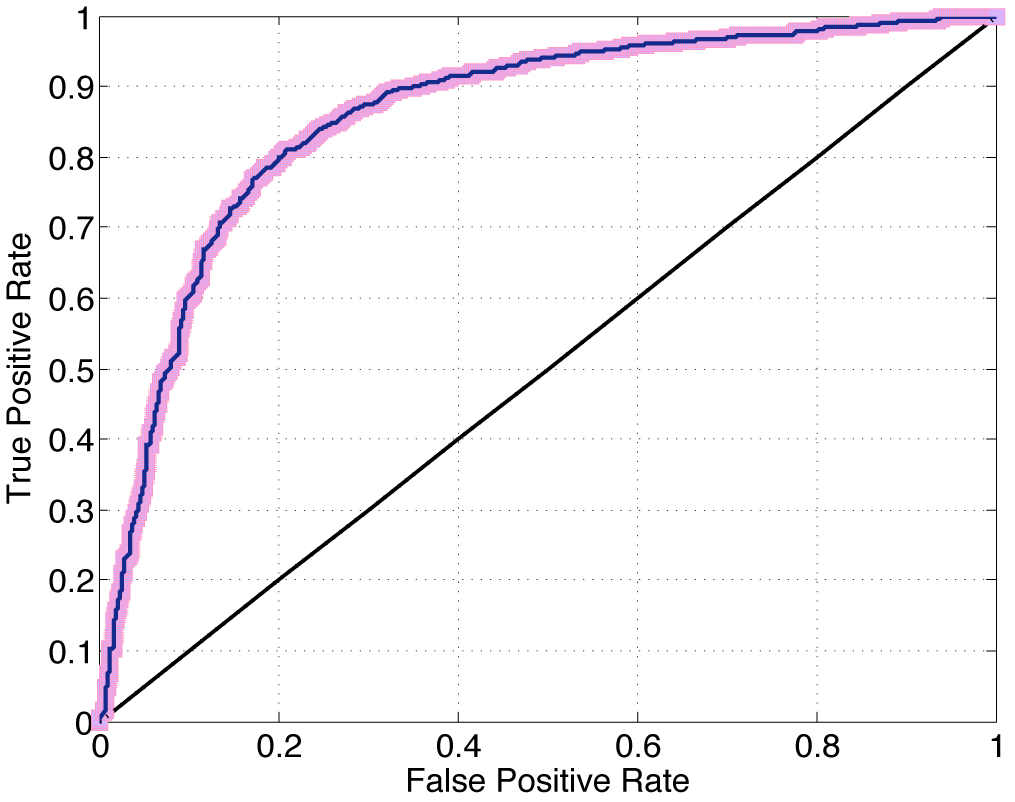
Discriminative identification of TrEn (‘robust-set’) versus random controls. We show the characteristic ROC curve using as input the most informative set of 31 motifs.

Next, we compare the results obtained using the ‘robust set’ of TrEn with the results achieved in the previous subsection by analysing the ‘all facets’ dataset. In particular, by aggregating the motif signatures per cell type/tissue from the ‘all facets’ dataset we observe that specific motifs are selected with high frequency across 112 cell types/tissues. Figure 4 shows the set of 31 informative motifs obtained by the ‘robust set’ (x-axis) and the selection frequency of every individual motif (y-axis) across 112 cell types/tissues from the ‘all facets’ dataset. We observe that six out of the 31 motifs are selected with more than 80% of times across different cell types/tissues. Notably, these motifs, namely CG, CGA, TCG, CGT, ACG and TA, almost always help maximizing discrimination performance. We also observe that two di-nucleotides appear very discriminative, CG and TA, and this finding has been also explored in experiments in Drosophila^20^ and reported by other independent studies^5^. From 31 reported motifs, 10 are tri-nucleotides rich in CG, whereas 19 are tetra-nucleotides again rich in CG.

**Figure 4:**
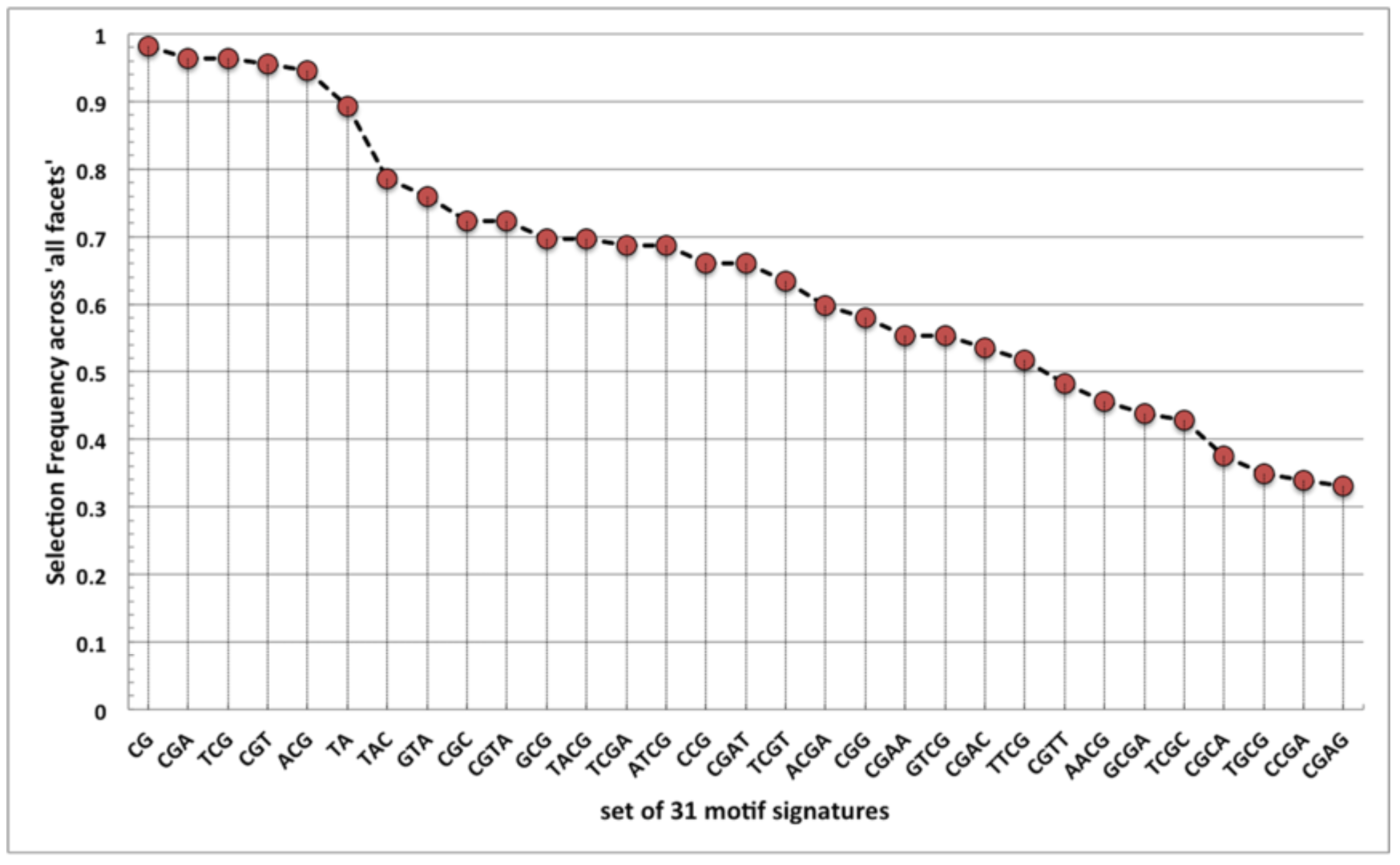
The most informative set of 31 motifs for discriminating the ‘robust set’ of TrEn versus random controls. The y-axis indicates the selection frequency of each motif obtained by analyzing the ‘all-facets’ dataset.

### Identification of DNA signatures of TrEn transcribed exclusively in only one cell type/tissue from FANTOM5

The hypothesis we investigate in this subsection is whether or not FANTOM5 enhancers expressed and transcribed exclusively in one cell type/tissue can be distinguished from exclusively transcribed enhancers in other cell types/tissues based on sequence characteristics. To explore this hypothesis, we apply TELS to identify motif signatures that discriminate effectively ‘exclusively transcribed’ enhancers from their corresponding ‘negative exclusively transcribed’ (see Materials and Methods for data description). The classification performance achieved across 96 cell types/tissues is presented in Supplementary Figure 2. Note that out of 112 cell types/tissues 16 have less than five exclusively transcribed enhancers and thus are filtered out. This particular discrimination problem is much more complex as represented by a drop in the classification performance compared to the results presented in the previous subsections. We show that ‘exclusively transcribed’ enhancers can be distinguished from ‘negative exclusively transcribed’ with an average PPV and GM of 65.23% and 65.02%, respectively. In some cell types/tissues (∼25 cases) the results are quite satisfactory with PPV greater than 80%, whereas in ∼40 cell types/tissues PPV ranges from 50% to 60%, which indicates that in addition to the identified DNA signatures, other mechanisms have strong influence on cell type/tissue-specificity. Figure 5 is in concordance with Figure 1 showing that the maximization of the classification performance is achieved using motif signatures of different sizes. However, the classification performance as indicated by PPV, is much lower compared to the case depicted in Figure 1.

**Figure 5:**
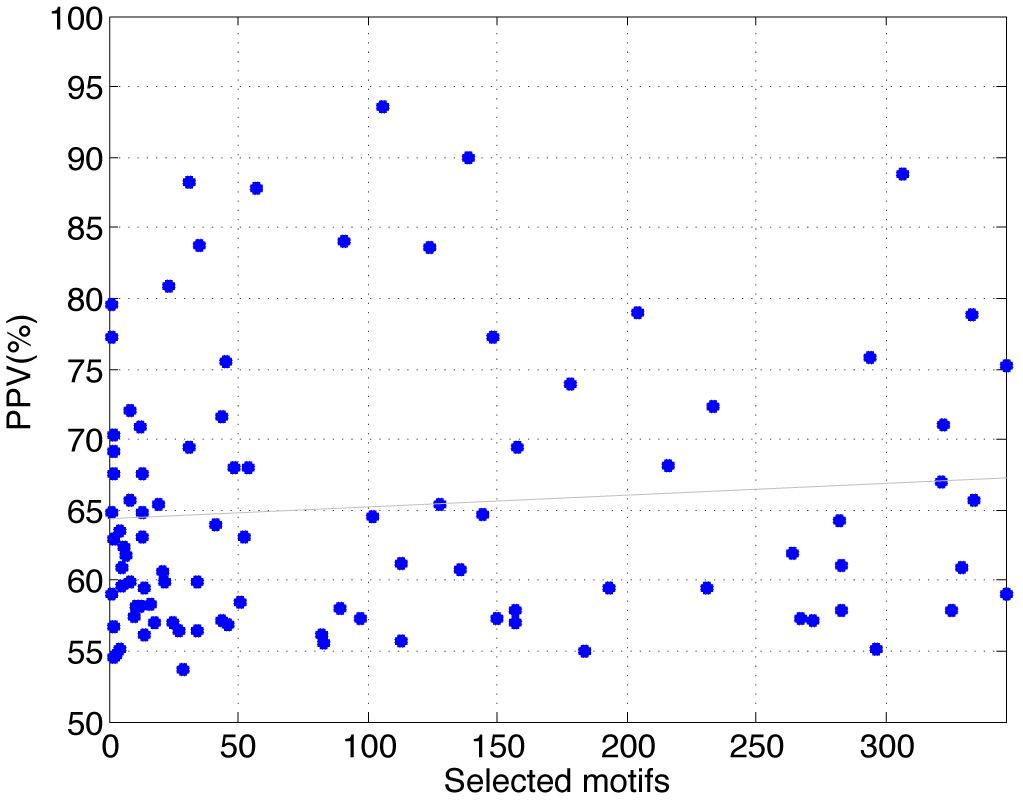
Number of selected motifs across 96 cell types/tissues versus the corresponding PPV (%) for identifying the ‘exclusively transcribed’ dataset from TrEn expressed in different cell type/tissues.

### Comparison to other sets of motif signatures

We compare the discriminative capabilities of motif signatures identified by TELS with motifs reported by other studies. These include: a) the results by Lee *et al.*^21^ who reported a set of 20 informative 6-mers that were used by linear Support Vector Machine (SVM) to distinguish chromatin-defined enhancers from random DNA sequences; b) the results by Yanez-Cuna *et al.*^20^ who performed STARR-seq experiments in Drosophila and reported motifs CA and GA, as well as the AP-1 motif among the most characteristic for enhancer activation; and c) the results of DEEP^35^, which utilized a set of 351 sequence characteristics under a complex ensemble model of 1000 SVMs to predict both transcribed and chromatin-defined enhancers on a genome-wide scale.

We note that comparing motif signatures identified by different computational approaches is not straightforward for several reasons. First, different methods are based on different enhancer and non-enhancer datasets. For example Lee *et al.* use enhancers defined by ChIP-seq, whereas Yanez-Cuna *et al.* use a quantitative experimental approach to measure enhancers activity in Drosophila. In addition, the enhancer’s sequence length has a great impact on the sequence composition and on the selection of motif signatures. We note that in our study, the TrEn length does not rely on ad-hoc windows of fixed length surrounding ChIP-seq peaks or STARR-seq peaks but derives from the TrEn length estimated in ref. (5). There are also differences in the selection of machine-learning models and tuning of model parameters (e.g., C parameter for SVM or number of SVMs in the ensemble).

Here, to make the comparison as fair as possible we use two independent classifiers, the KNearest Neighbours (KNN) and Bagged Decision Trees (BDT). Both classification algorithms are not used for training models by any of the competitor methods. KNN and BDT are implemented in Matlab and run in a default setting for all experiments (K=3 for KNN and B=20 for Decision Trees). To assess the classification performance we repeat the learning process 100 times and in every run we split the data randomly into training (60%) and testing (40%) sets.

We visualize the distribution of PPV and GM of 100 runs using box diagrams (Figure 6). It is apparent that the set of 31 motifs identified by TELS discriminates much more accurately the ‘robust set’ of TrEn compared to motifs identified by other studies. This outcome is consistent across two different classification algorithms. In fact, the results presented here indicate that the motif signatures reported by TELS are characteristic to the category of CAGE-defined enhancers whereas the motifs reported by the competitor methods are tailored to chromatin-defined enhancers or enhancers expressed in other species.

**Figure 6:**
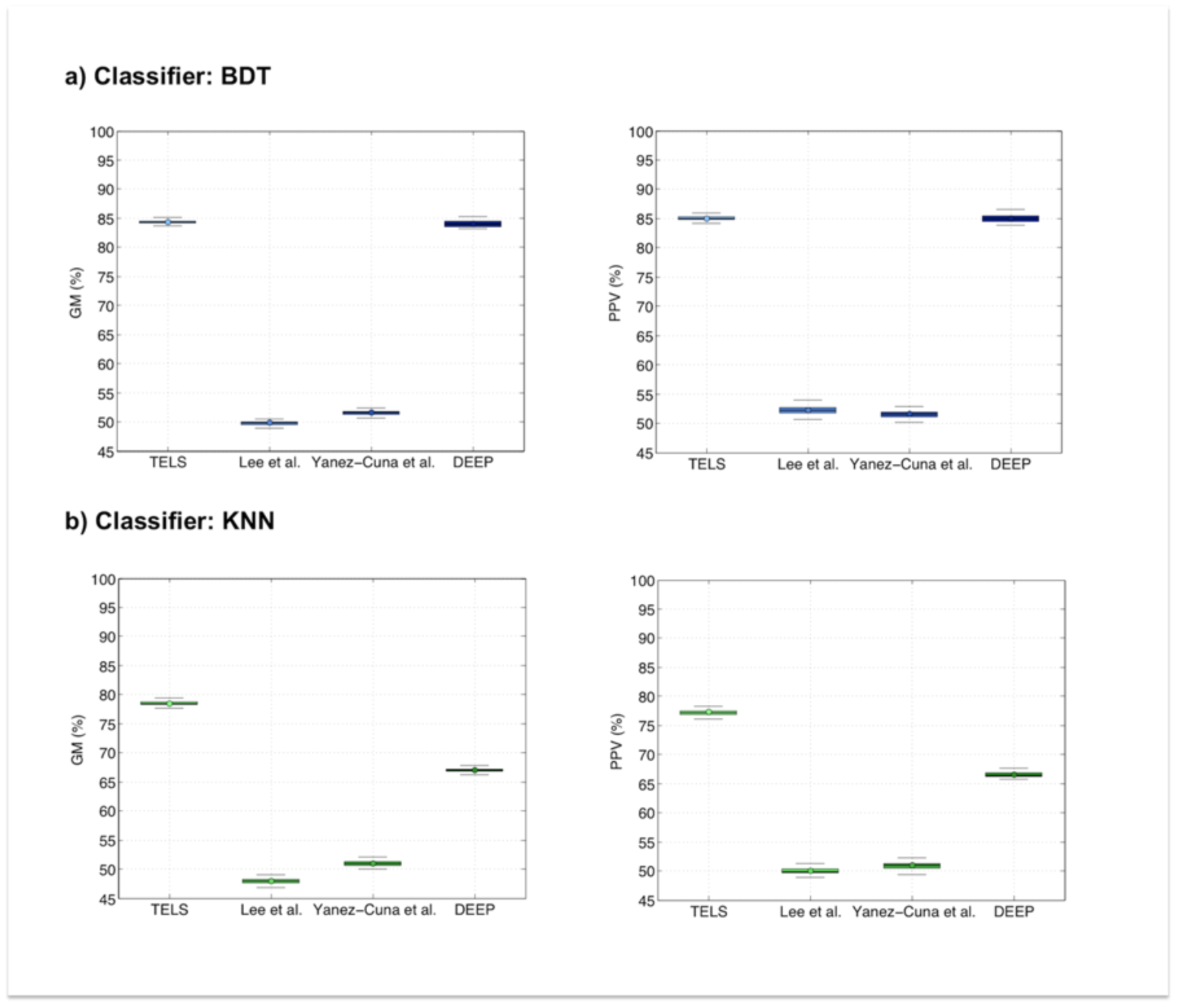
Comparison analysis of motifs identified by alternative approaches: (a) Classification performance in terms of GM (%) and PPV (%) using BDT; (b) Classification performance in terms of GM (%) and PPV (%) using KNN.

From the technical point of view, TELS achieves comparable classification performance using three independent classification methods namely LR, BDT, and KNN. This indicates that the findings are not biased to one particular classification model (i.e., we used LR during the motif selection). Moreover, our results indicate that the classification algorithm used for assessing the motifs’ importance resulted in no bias in the motif selection process

### Validation using ENCODE enhancers

As an additional validation process we test the discriminative capabilities of the identified motif signatures on completely independent datasets. For this purpose, we utilize chromatin-defined enhancers reported by the ENCODE consortium^36^ that do not overlap with the CAGE-defined enhancers from the FANTOM5 Atlas^5^. In this way, we perform ‘blind’ validation, but most importantly, we investigate the degree to which different sequence properties of transcribed enhancers can be utilized to discriminate effectively enhancers identified by chromatin characteristics.

We use the list of ‘strong’ enhancers^36^ from six cell-lines (see Materials and Methods for data description). As input variables, we use the set of 31 motif signatures derived from the ‘robust set’ of TrEn. For classification we use KNN and BDT algorithms under the default setting (K=3 for KNN and B=20 for Decision Trees). For assessing the classification performance we measure the average PPV and GM of 100 runs, where in each run we split the data randomly into training (60%) and testing (40%) sets.

The distribution of PPV and GM using box diagrams is presented in Figure 7.Our results demonstrate, that the set of 31 motif signatures discriminates with high accuracy chromatin-defined enhancers from random counterparts. This finding supports the hypothesis that TrEn and chromatin-defined enhancers have similar DNA specificities and thus similar sequence motifs are effective in both enhancer types. Biologically this might also indicate that many of the ‘strong’ enhancers defined by ChIP-seq are transcribed and/or that there are some common DNA sequence characteristics for all poised and active enhancers. More importantly, the results of this section re-confirm that TELS can be used to decipher systematically the motif signatures of distal regulatory elements.

**Figure 7:**
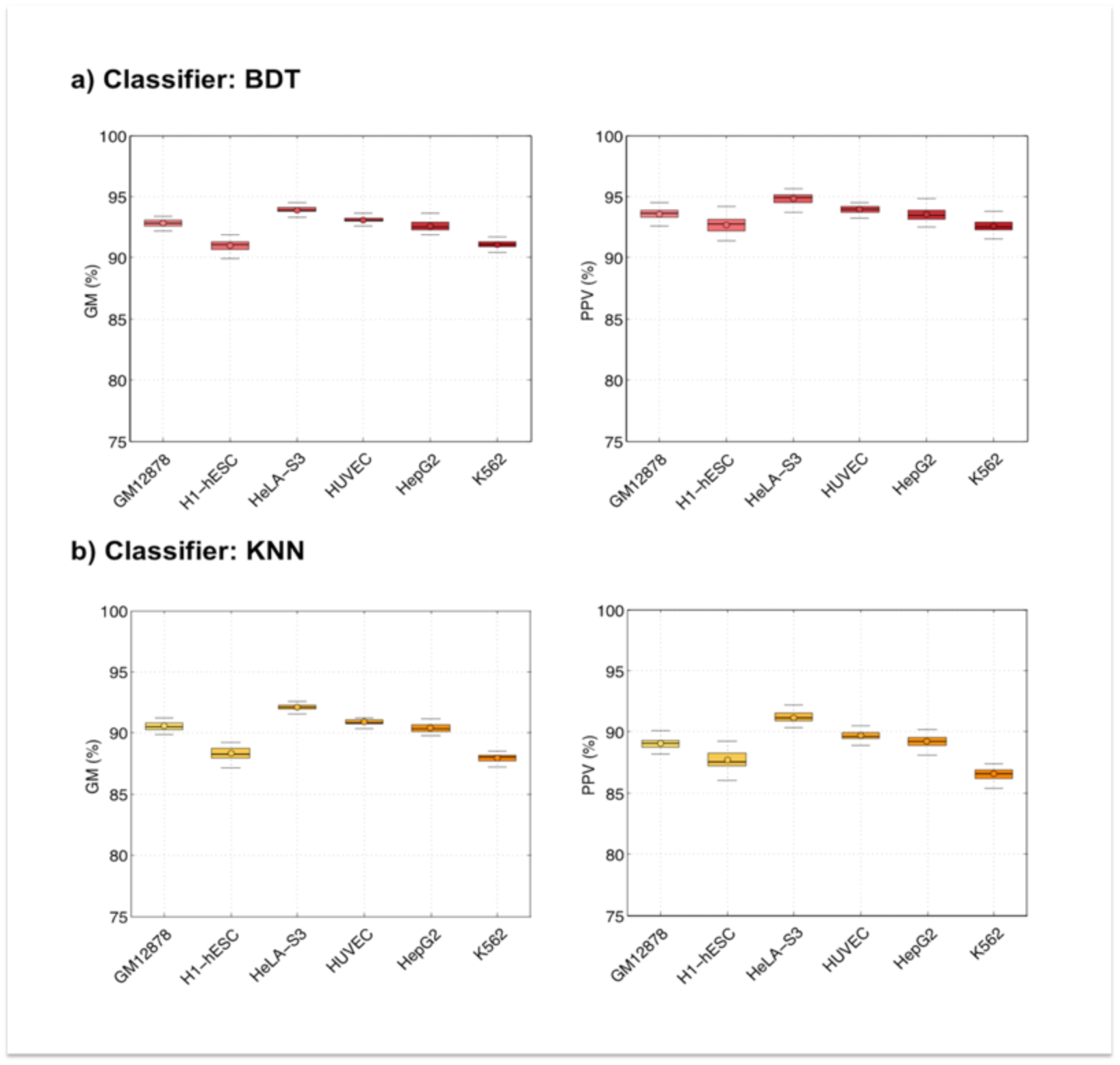
Classification performance using the set of 31 motif signatures tested on chromatin-defined enhancers from ENCODE: (a) Performance in terms of GM (%) and PPV (%) using BDT; (b) Performance in terms of GM (%) and PPV (%) using KNN.

## MATERIALS AND METHODS

### Data availability

The primary datasets included in our study derive from the FANTOM5 Atlas of TrEn^5^. Using a large number of primary cells and tissues, that study identified bi-directional TrEn via CAGE experiments. All enhancer samples were obtained from the Atlas webpage (http://enhancer.binf.ku.dk/presets) on November 2016. Details about the TrEn identification pipeline from CAGE and other information about the primary data can be found in ref. (5).

For validating further our findings, we use the list of ‘strong’ enhancers reported by the ENCODE integrative annotation^36^. Details about the ‘strong’ enhancer identification process can be found in ref. (36). We focus on six ENCODE cell-lines namely:GM12878, H1-hESC, K562, HeLa-S3, HepG2 and HUVEC. From the cell-line specific lists of ‘strong’ enhancers we consider only the sequences that do not overlap with CAGE-defined enhancers from the FANTOM5 TrEn Atlas^5^. This guarantees that the ENCODE data we use for testing are completely independent from the FANTOM5 data we use for identifying motif signatures.

All datasets generated in this study as well as TELS source codes for reproducing the results are publicly available at http://www.cbrc.kaust.edu.sa/TELS under an Educational Community Open Source Licence. In the Supplementary Material we also provide a simple flowchart of TELS (Supplementary Figure 3) as well additional information about Materials and Methods.

### Definition of positive and negative datasets

To identify motif signatures of TrEn and develop discrimination models, we use the following three datasets that are considered ‘positive’ data:

a. ‘all facets’ (http://enhancer.binf.ku.dk/presets/facet_expressed_enhancers.tgz): This contains enhancers transcribed in all FANTOM5 facets from 112 cell-types and tissues (i.e., 112 TrEn sets). This dataset contains 197,373 genomic sequences including duplicates since some TrEn are expressed and transcribed in more than one cell type/tissue.
b. ‘robust set’ (http://enhancer.binf.ku.dk/presets/robust_enhancers.bed): This set contains the enhancers transcribed at a significant expression level in at least one FANTOM5 primary cell/tissue. This dataset contains 38,554 genomic sequences.
c. ‘exclusively transcribed’: We denote as exclusively transcribed enhancers those TrEn that are transcribed in only one FANTOM5 cell type/tissue. We generate a list of exclusively transcribed enhancers for every cell type/tissue from the ‘all facets’ dataset described above. Basophil and granulocyte cell types have no exclusively transcribed enhancers. We also exclude from ‘exclusively transcribed’ dataset cell types/tissues with less than five exclusively transcribed enhancers. This results in 96 out of 112 potential datasets (one set per cell type/tissue).

Generating ‘negative’ data for the previously described ‘positive’ datasets without experimental validation (i.e., MPRA or STARR-seq) is a difficult task since it is unclear based only on computational consideration to infer whether or not a particular DNA sequence has enhancer activity in a cell type/tissue. To mitigate this problem we considered two alternative approaches: i) based on artificially generated sequences; and ii) using ‘real’ TrEn expressed in different cell types/tissues than the one of interest.

The rational behind (i) derives from studies showing that mutations in enhancer sequences disrupts the enhancer’s activity^12,16,18,19^. Thus, for every TrEn in the ‘all facets’ dataset as well as for the ‘robust set’, we mutate randomly 10% of the genomic positions in every TrEn sequence. We decided to mutate 10% of the positions and not less to have a better proxy of the non-enhancers. In this way, from every ‘real’ TrEn we generate a random negative sample that has exactly the same sequence length but with shuffled DNA composition^22^. We generate in total 197,373 negative controls across 112 cell types/tissues named as ‘all facets random controls’ and 38,554 negative controls named as ‘robust set random control’ respectively. To note, that throughout the previous data generation process, we make sure that none of the randomly generated sequences belong to the superset of TrEn identified by FANTOM5. In (ii) we follow the ‘one vs. all’ paradigm and for every cell type/tissue specific ‘exclusively transcribed’ enhancer set, we generate a negative set that contains exclusively transcribed enhancers from all other cell types/tissues but not from the one of interest. This process resulted to 96 negative sets and such dataset is denoted as ‘negative exclusively transcribed’.

For all ‘positive’ and ‘negative’ datasets, we use the enhancer genomic coordinates provided in the bed format to generate the actual DNA sequences in the fasta format using SAMtools^23^ and the UCSC reference genome (assembly version hg19).

### DNA sequence encoding

To encode the input datasets for further use by TELS, we transform all ‘positive’ and ‘negative’ data samples into numerical vectors. In TELS, we focus on small sequence motifs and not on preselected TFBSs from PWM or ChIP-seq. In this way, we consider the intrinsic DNA properties of TrEn and we complement similar studies that focus on known motifs usually of length of 6 or 10 bp. The deployed vectors contain 346 variables (equally denoted in this manuscript as sequence motifs or motifs, or features) that describe the enhancers’ genomic specificity. These variables are grouped into four categories: a) four single nucleotide frequencies; b) six aggregate frequencies of two nucleotides (e.g., A+C); c) 16 dinucleotide frequencies; d) 64 trinucleotide frequencies; and e) 256 tetranucleotide frequencies. To avoid any bias introduced by the length of the sequences, we normalize all values of the vectors by the sequence length.

### TELS implementation

TELS works in two phases. In phase (1), TELS identifies candidate combinations of sequence motifs that characterize the class of interest. In phase (2) for every candidate combination of motifs, TELS assesses its significance by measuring the classification performance for discriminating ‘positive’ from ‘negative’ data. The objective of TELS is to select the combination of motifs that maximize separation between ‘positive’ and ‘negative’ data. Typically, determining the relative importance of a set of predictor variables via computational techniques may be used to associate differences between the considered data classes. This information can be further utilized to identify sequence characteristics that are predictive of TrEn cell type/tissue specific activities.

In phase (1), to identify candidate combinations of motifs TELS uses filtering feature selection (FS) techniques. The FS problem in bioinformatics is very well studied^25-33^ and it is well documented that FS is a strongly ‘data-dependent’ process. TELS first ranks the 346 individual variables using the Gini-index based FS. We decided to use Gini-index after comparison with two other state-of-the-art algorithms for FS, namely minimum redundancy maximum relevance criterion (mRMR)^34^ and Fisher’s test-based FS^33^ (Supplementary Material). We used Gini-index implementation from the Feature Selection Toolbox (FEAST) in Matlab R2014b.As importantly, feature ranking by filtering methods ignores interaction with classification models. Thus, features are considered ‘separately’, which may lead to suboptimal classification performance. In other words, from a pool of 346 ranked variables based on their significance assessed by Gini-index, is not clear which combination characterizes in an optimized manner the class of interest.

To mitigate this problem in phase (2), we follow a greedy approach, and assess the significance of all possible combinations of ranked features (starting with the top-1, top-2, top-3 and up to top-K, where K is 346, the total number of variables we use) by measuring the classification performance of every candidate combination. Our objective is to select the combination of motifs that based on MCC minimizes the classification error (i.e., maximizes classification performance). For this task TELS utilizes the LR classifier. LR is a simple linear classification method, which runs fast and avoids extensive optimization of model parameters that frequently leads to poor performance on unseen data^24^. The implementation is made in Matlab R2014b using built-in functions for LR (‘glmfit’ function with the default setting).

In one classification run with LR and one candidate motif set, we randomly split the ‘positive’ and ‘negative’ data into testing and training sets. To achieve better generalization on unseen data, we use 20% of the total size of ‘positive’ and 20% of the total size of ‘negative’ samples for training and the remaining 80% from both is kept for testing. We finally report the average classification performance of 300 runs when for each run the above mentioned random split of data is performed. Consequently, we estimate the average classification performance of 300 runs for all candidate combinations of top-ranked motifs (i.e., 346) and this guarantees an equitable selection of the combination that maximizes classification performance. We consider Geometric mean of Sensitivity and Specificity (GM), Positive Predictive Value (PPV), and Mathews Correlation Coefficient (MCC) as representative classification performance metrics. All performance metric formulas can be found in the Supplementary Material.

## CONCLUSION

In this study we proposed TELS, a novel machine-learning algorithm for identifying predictive short motif signatures of TrEn. We tested TELS using CAGE-defined enhancers from FANTOM5. This allowed us to compile a comprehensive catalog of cell type/tissue specific combinations of short motif signatures that maximize discrimination using different negative datasets. Specifically, our analysis reported combinations of motifs that allowed us to discriminate effectively TrEn from random controls, a proxy of non-enhancer activity. We also reported combinations of motifs that maximize classification performance of exclusively transcribed enhancers in one cell type/tissue from enhancers transcribed exclusively in all other cell types/tissues. In addition, by analysing the so-called ‘robust set’ of TrEn we were capable of finding 31 motif signatures predictive of TrEn broad activity. We also showed that this set is quite universal and can discriminate with high performance chromatin-defined enhancers from six different ENCODE cell-lines. However, we note that findings obtained by any computational method require further targeted wet-lab experiments even when supported by statistical evidence.

Nonetheless, within the present bioinformatics method, many improvements are possible:

a. Replacing LR with a non-linear classifier such as DBT might improve further the already high discrimination performance;
b. Performing the same analysis on TrEn data obtained by single-cell analysis, if available by FANTOM, will eliminate potential biases caused by cell population heterogeneity and may lead to more fine-grained results about the enhancer genomic landscape;
c. Applying the same analysis to CAGE-defined promoters from FANTOM5 will answer equally important questions about promoters’ sequence characteristics ‘encrypted’ within their genomic sequence;
d. Stratifying TrEn data by their expression levels similar to ref. (9), and analysing with TELS the sequence composition of different expression subclasses, may complement the findings presented in this study.

## ACKNOWLEDGEMENTS

The research reported in this manuscript was supported by the base funding grant No. BAS/1/1606-01-01 of the King Abdullah University of Science and Technology (KAUST) to VBB.

## CONTRIBUTIONS

D.K and V.B.B conceived the project, analysed the data and wrote the manuscript text. H.A and N.Z performed the experiments. All authors read and approved the final manuscript.

